# A global signature of microbiome resilience to climate stress

**DOI:** 10.64898/2026.09.30.755659

**Authors:** Amanda H Rawstern, Alma L Reyes, Vicki W Li, Joshua C Fowler, Eleanor M Hay, Damian J Hernandez, Anne M Katula, Ahmed Gebriel, Michelle E Afkhami

## Abstract

As climate change accelerates, the microbial communities sustaining organisms and ecosystems worldwide face an uncertain future, with some proving resilient to stress while others collapse, yet no generalizable framework exists to distinguish the two beforehand. Using amplicon sequencing data from ∼29,000 microbiomes, we constructed over 440 matched network pairs spanning three of the most pressing stressors: drought, warming, and salinity. We show that keystone microbes with high connectivity, functional redundancy, and metabolic flexibility, are disproportionately likely to survive stress, and that baseline network and functional properties predict community-level structural and functional stability even as taxonomic composition shifts stochastically. These relationships held across the diverse hosts, ecosystems, and regions we tested. Because these properties can be derived from standard, low-cost sequencing collected before disturbance occurs, our findings turn baseline microbiome data into an early-warning system for identifying which communities are most vulnerable to climate stress.

## Introduction

Climate change is accelerating at an unprecedented rate, driving widespread shifts in temperature, precipitation, and salinity^1^ that are fundamentally destabilizing natural and agricultural ecosystems worldwide^2^. At the foundation of these ecosystems lie microbial communities, the unseen drivers of nutrient cycling, carbon sequestration, and greenhouse gas regulation that sustain life on Earth^3–4^. Beyond their roles in global biogeochemical cycles, microbes also provide direct climate protection to plants and animals, producing osmolytes that aid water retention, reducing salt toxicity, and forming biofilms that shield against UV damage and thermal stress^5–6^. Despite their outsized role in ecosystem function, microbiomes remain vastly understudied relative to macroorganisms in the context of climate vulnerability, representing a critical blind spot in our understanding of how ecosystems will respond to intensifying stress^7^.

Yet microbiomes themselves are highly vulnerable to climate stress, with intensifying abiotic stress driving dysbiosis^8^, disrupting host-microbiome mutualisms^6^, and causing the local extinction of keystone taxa whose loss can trigger cascading functional collapse^9^. These disruptions carry profound consequences for both natural and managed ecosystems. Microbiomes govern plant productivity, soil fertility, nutrient availability, and biodiversity that terrestrial and marine ecosystems depend upon, making their stability essential for sustaining both food security and ecosystem integrity worldwide^10–11^. However, the extent to which microbiomes are destabilized by climate stress varies widely, with some communities proving remarkably resilient while others undergo rapid structural and functional collapse^12^. Despite decades of calls for integrating microbiomes into biodiversity monitoring and conservation frameworks^7,13^, no generalizable predictive framework exists that allows researchers and land managers to assess climate vulnerability using standard, pre-disturbance microbiome data collected from their own systems.

Ecological theory posits that the structural and functional organization of microbial communities should underpin their capacity to resist and recover from disturbance^6,13–14^. Among network structural properties modularity compartmentalizes taxa into discrete interaction groups, limiting the spread of perturbation across the broader community^15–16^. Cohesion reflects the balance of negative and positive microbial associations, where communities dominated by negative interactions prevent over-reliance between taxa and reduce the risk of cascading co-extinction, while communities approaching balanced cohesion retain sufficient cooperative interactions to maintain community function and host benefits^14,17–19^. Functional redundancy, the capacity of multiple taxa to perform equivalent ecological roles, provides an additional buffer against climate-driven diversity loss by ensuring that critical functions persist even when individual taxa are lost^20–21^. However, this buffering capacity is not uniform across all functions, broad functions performed by a wide range of taxa tend to exhibit higher redundancy than narrow, specialized functions restricted to relatively few taxa^22^. Keystone taxa exert disproportionate influence on community diversity and function regardless of their abundance, and their retention under stress is critical for preserving these narrow, specialized functions that drive community recovery^9,23–24^. Despite the theoretical promise of these properties, their utility as generalizable predictors of climate resilience remains largely untested. The majority of existing studies examine single systems with few network pairs, and inconsistent sampling designs where some studies capture only pre-disturbance or only post-disturbance communities make it impossible to determine whether these properties reliably predict stability across diverse ecosystems and stressors on a global scale^13,15^.

Here we address this gap by conducting novel network and functional analyses on over 29,000 individual microbiomes from publicly available raw amplicon sequencing data. We developed a standardized pipeline encompassing co-occurrence network construction and functional prediction to generate over 440 matched ambient-stressed network pairs. These networks spanned host-associated and free-living communities across terrestrial and marine ecosystems. We focused specifically on drought, warming, and salinity, three of the most pervasive and ecologically consequential climate stressors driving biodiversity loss and ecosystem destabilization worldwide. We demonstrate that easily obtainable baseline microbiome community attributes, including modularity, negative to positive cohesion, functional redundancy, and mean keystone centrality, are strong and generalizable predictors of structural and functional stability across diverse stressors and ecosystems. Critically, by testing stability across three ecological dimensions, structural, functional, and compositional, we show that overall community stability remains predictable even when accounting for the stochastic nature of compositional turnover, suggesting these properties capture the core determinants of microbiome resilience under climate stress. Together these models provide actionable predictive tools that scientists can apply to their own baseline microbiome data to forecast keystone retention and overall community stability, enabling identification of the ecosystems most vulnerable to climate-driven microbiome collapse before stress events occur.

## Results

### Global amplicon analysis reveals generalizable predictors of microbiome climate resilience

To investigate how baseline microbial network properties predict community stability under climate stress, we compiled an extensive global dataset of microbiomes derived from publicly available amplicon sequencing data spanning three of the most prevalent climate stressors affecting terrestrial and aquatic ecosystems. Using fast, low-cost methods including amplicon processing, FastSpar co-occurrence network construction, and PICRUSt2 functional prediction we generated 441 network pairs (882 total networks). Each pair represented matched microbiomes originating from the same environment and sample source under ambient and climate-stressed conditions, drawn from 110 studies encompassing over 29,000 microbiome samples (Supplementary Table 1). Studies spanned drought (n = 243 network pairs), warming (n = 123), and salinity (n = 75) stressors (Fig. 1a), and included both prokaryotic (n = 315) and fungal (n = 126) communities from host-associated (n = 251) and free-living (n = 190) environments distributed globally (Fig. 1b). Together, these data demonstrate that widely accessible sequencing and computational methods are sufficient to capture the structural and functional microbial community properties needed to predict microbiome stability across diverse stressors, host types, and ecosystems.

**Fig. 1:**
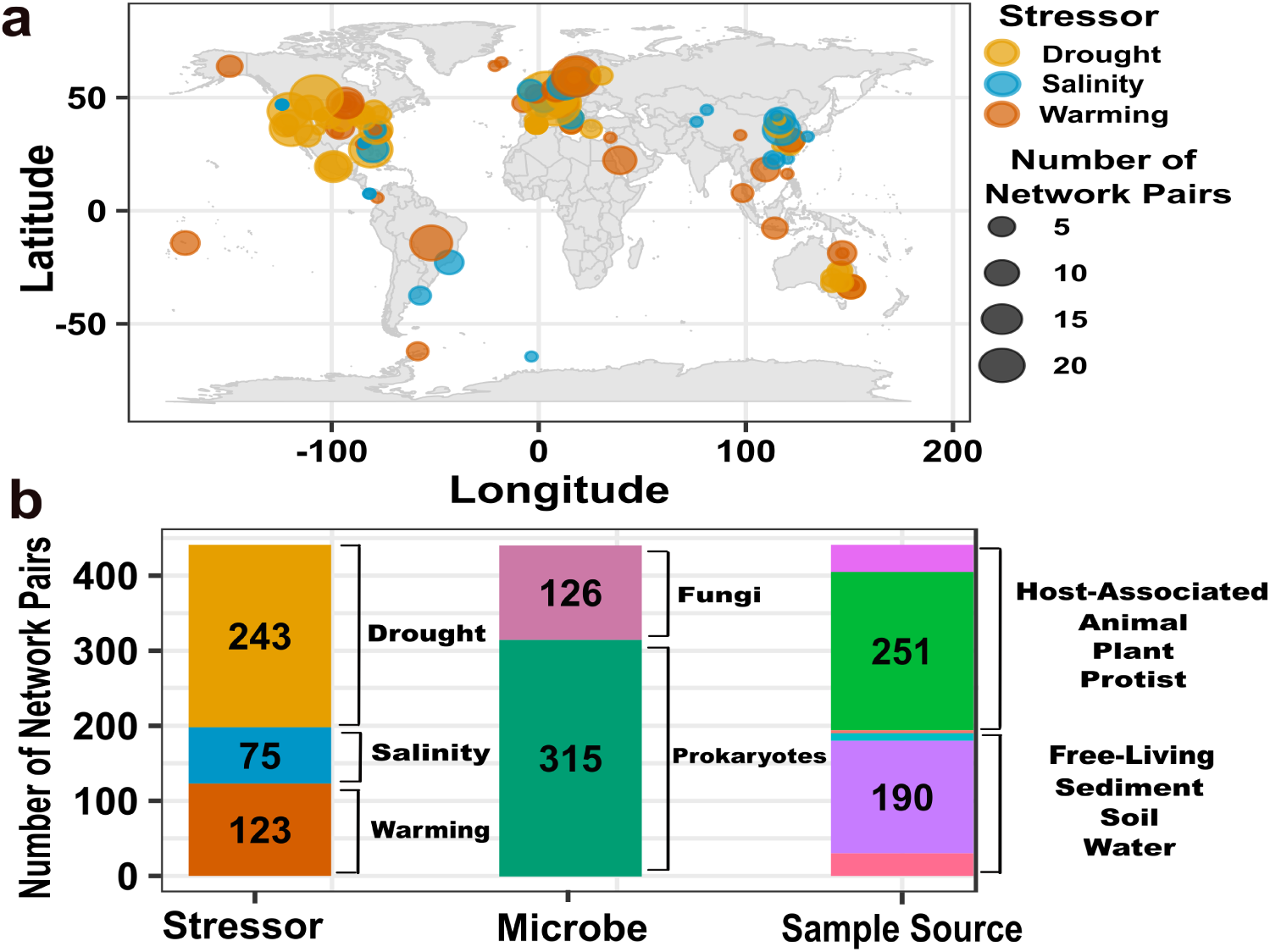
Global microbiome dataset spans a diversity of hosts and environments. (a) Geographic distribution of network pairs across studies, with circle size indicating the number of network pairs per location and color indicating stressor type (n = 441 network pairs encompassing n = 29,005 microbiome samples from n = 110 studies). (b) Breakdown of the network pairs by stressor (drought, n = 243; warming, n = 123; salinity, n = 75), microbe type (prokaryotes, n = 315; fungi, n = 126), and sample source (host-associated, n = 251; free-living, n = 190).

### Functional depth and network connectivity distinguish retained from lost keystone microbes under climate stress

We identified keystone taxa across all 441 network pairs and tracked their retention or loss following climate stress events. Using indicator analysis we identified 5 retention-associated enzymatic functions significantly enriched in retained keystones (Fig. 2a), including ammonia monooxygenase (EC:1.14.99.39), which supports nitrogen cycling under shifting nutrient conditions; particulate methane monooxygenase (EC:1.14.18.3), which enables energy acquisition under fluctuating redox conditions; 3-hydroxypropionyl-CoA dehydratase (EC:4.2.1.116), which supports autotrophic CO2 fixation independent of organic carbon availability; rRNA small subunit pseudouridine methyltransferase Nep1 (EC:2.1.1.260), associated with translational stress responses and cellular resilience; and 3-carboxy-cis,cis-muconate cycloisomerase (EC:5.5.1.2), which enables degradation of aromatic compounds released during stress-induced cell death. Together these functions suggest that retained keystones share a common capacity for metabolic flexibility under the resource-limited and chemically altered conditions characteristic of climate stress events. Consistent with this, retained keystones exhibited significantly greater functional redundancy, reflecting a greater ability to perform multiple functions within the same ecological category, than lost keystones (z = 14.26, p < 0.001; Fig. 2b, Extended Data Fig. 1a, Supplementary Table 2a), indicating that redundancy protects against the loss of critical functions under stress. Among keystone taxa, those with the highest network centrality were significantly more likely to be retained following stress (z = 16.21, p < 0.001; Extended Data Fig. 1b), suggesting that keystones interacting with a greater number of community members may be buffered against loss through a broader network of biotic interactions.

**Fig. 2:**
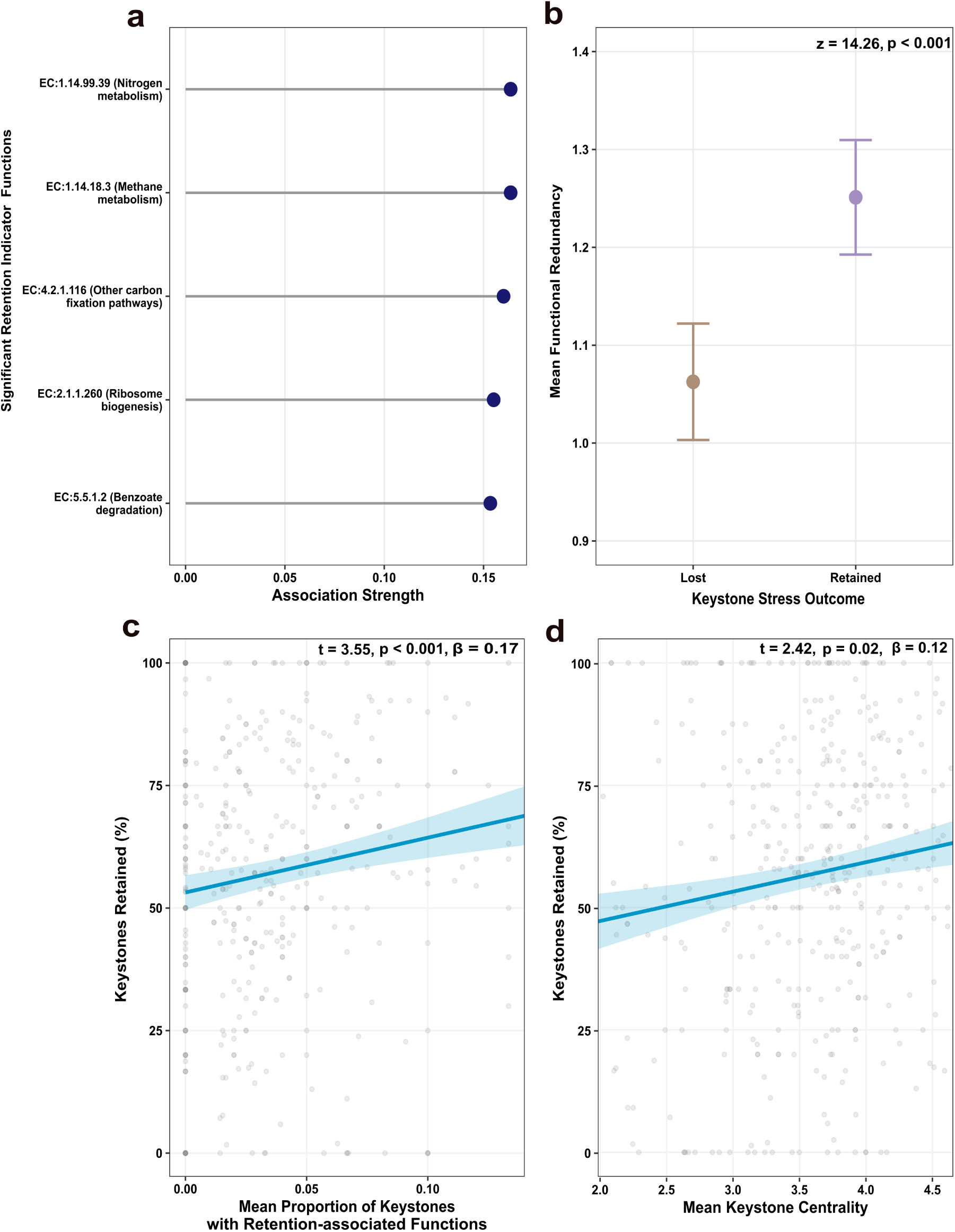
Keystone taxa functional profiles, functional redundancy, and network centrality predict whether keystone taxa is retained or lost after stress. (a) Indicator functions with KEGG annotations significantly associated with keystone retention across stress events with corresponding association strength (i.e. effect size of IndVal analysis). (b) Retained keystones exhibit higher functional redundancy than lost keystones (statistics given for logistic regression model). (c) Communities with higher proportions of keystones carrying retention-associated functions and (d) higher mean keystone centrality retain a greater percentage of keystones under stress (statistics given for linear mixed model with study as a random effect). Points and lines are mean values while error bars and shading represent 95% CIs.

These individual-level attributes translated into consistent patterns of collective keystone persistence. Communities with a higher proportion of keystones carrying retention-associated functions maintained significantly more keystones following stress (t = 3.55, p < 0.001, β = 0.17; Fig. 2c; Supplementary Table 2b), as did communities with higher mean keystone centrality (t = 2.42, p = 0.02, β = 0.12; Fig. 2d; Supplementary Table 2b), suggesting that both the functional repertoire and connectivity of the keystone community as a whole determine the retention of key taxa during climate stress. To further evaluate how well these attributes forecast individual and community keystone survival, we applied a Boruta-confirmed random forest model, which achieved strong classification performance (AUC = 0.775, accuracy = 0.713; Supplementary Table 2c). Keystone centrality emerged as the dominant predictor of survival, followed by functional redundancy and stressor type (Extended Data Fig. 2, Extended Data Fig. 3), confirming that a keystone’s functional depth and network position are the primary determinants of whether it survives climate stress, independent of stressor identity. Given the outsized roles keystone taxa play in community assembly, maintenance, and functional specialization, their persistence under stress is expected to have disproportionate consequences for overall community resilience.

### Baseline network and functional properties shield communities from structural and functional collapse despite stochastic compositional turnover

Linear mixed models with study as a random effect revealed that baseline network and functional properties significantly predicted all three dimensions of microbiome shift experienced during climate stress: structural (percent change in network topology), functional (percent change in community functional capacity), and compositional (percent change in taxa composition; Fig. 3, Fig. 4, Supplementary Table 3). Structural metrics encompassed network cohesion (ratio of negative to positive interactions), modularity (the degree of compartmentalization), and network richness (number of taxon nodes). Functional metrics comprised broad functional redundancy (mean number of functions per taxon across all functional categories) and narrow functional redundancy (mean number of functions per taxon within keystone functional categories). Building on the keystone persistence attributes identified in the preceding analysis, mean community keystone centrality and the proportion of keystones carrying retention-associated functions were also incorporated as community-level predictors of broader microbiome stability.

**Fig. 3:**
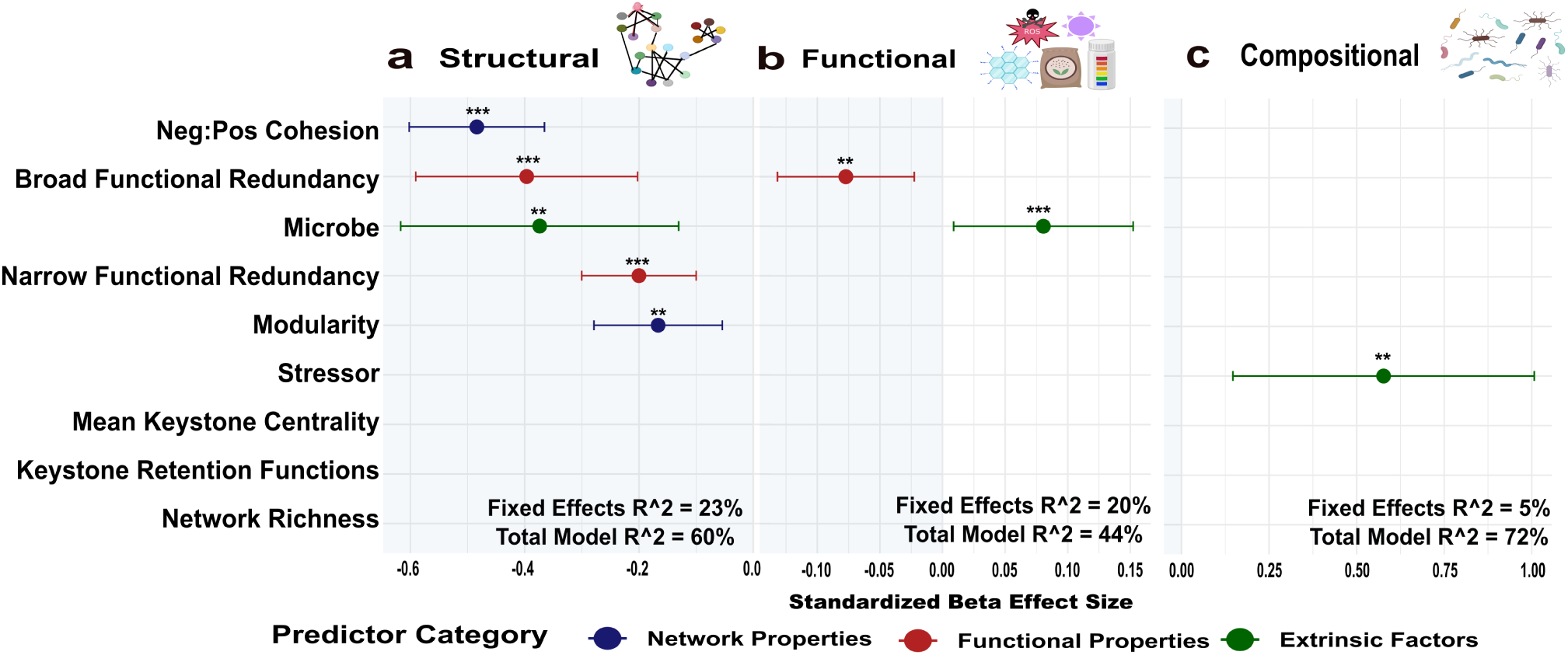
Baseline network properties predict microbial community stability under climate stress. Baseline properties predicting the (a) structural, (b) functional, and (c) compositional microbiome shift experienced during stress. Points denote effect size while error bars indicate 95% CIs. Points are colored by predictor category: network properties (blue), functional properties (red), and extrinsic factors (i.e. stressor and microbe type) (green). Statistics included are fixed and total model R² values and linear mixed model significances are indicated as **p < 0.01, ***p < 0.001.

**Fig. 4:**
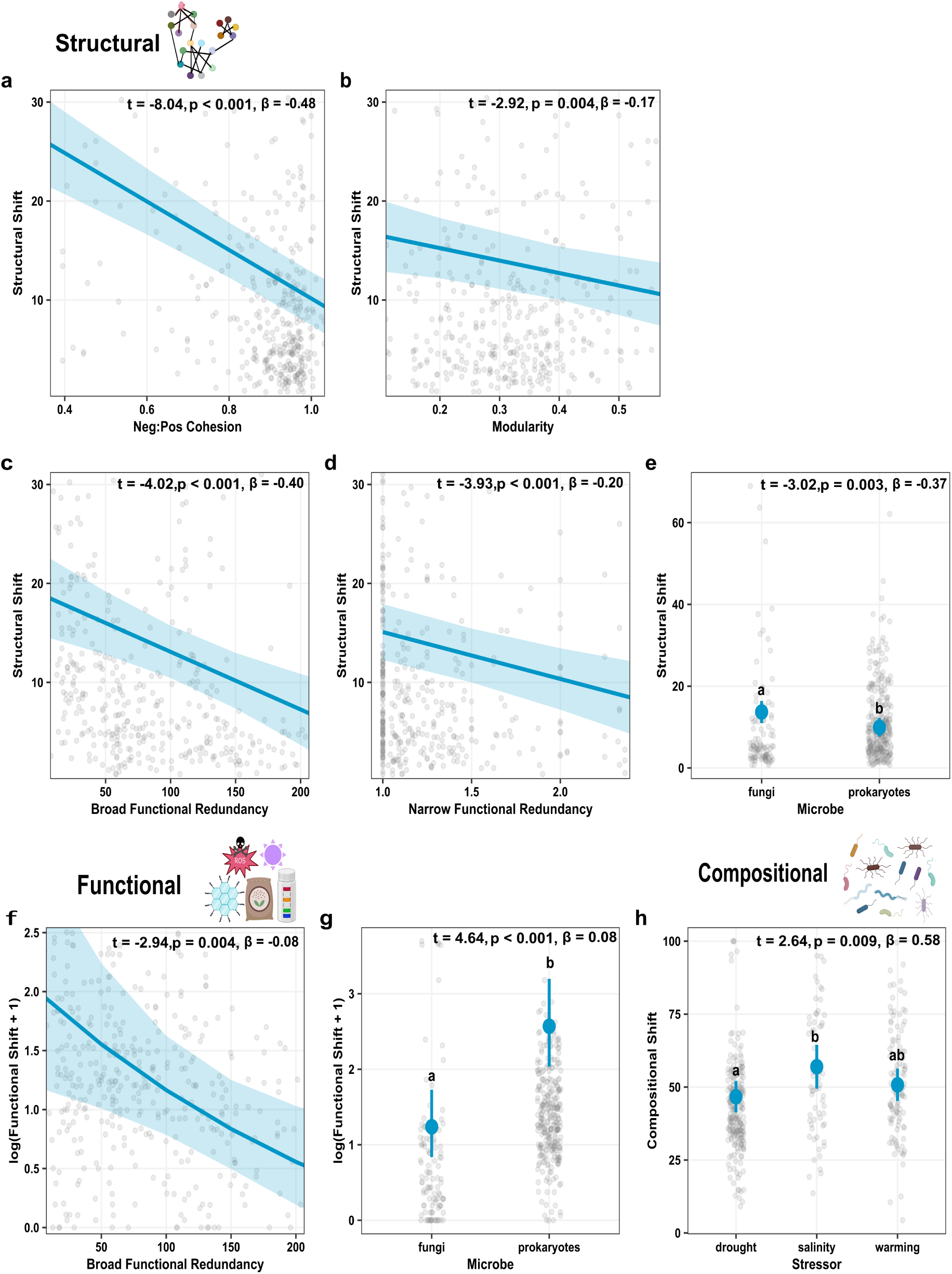
Partial effects of significant baseline predictors on structural, functional, and compositional shift. (a–d, f) Scatter plots show partial regression relationships between continuous predictors and stability dimensions while (e, g, h) dot plots show adjusted means for categorical predictors with ± 95% CI delineated for each plot. Mixed linear model t-values, standardized beta coefficients, and significance levels statistics are indicated in each panel. Panels are organized by baseline factors that significantly predict (a-e) structural, (f-g) functional, and (h) compositional shift experienced during climate stress.

**Fig. 5:**
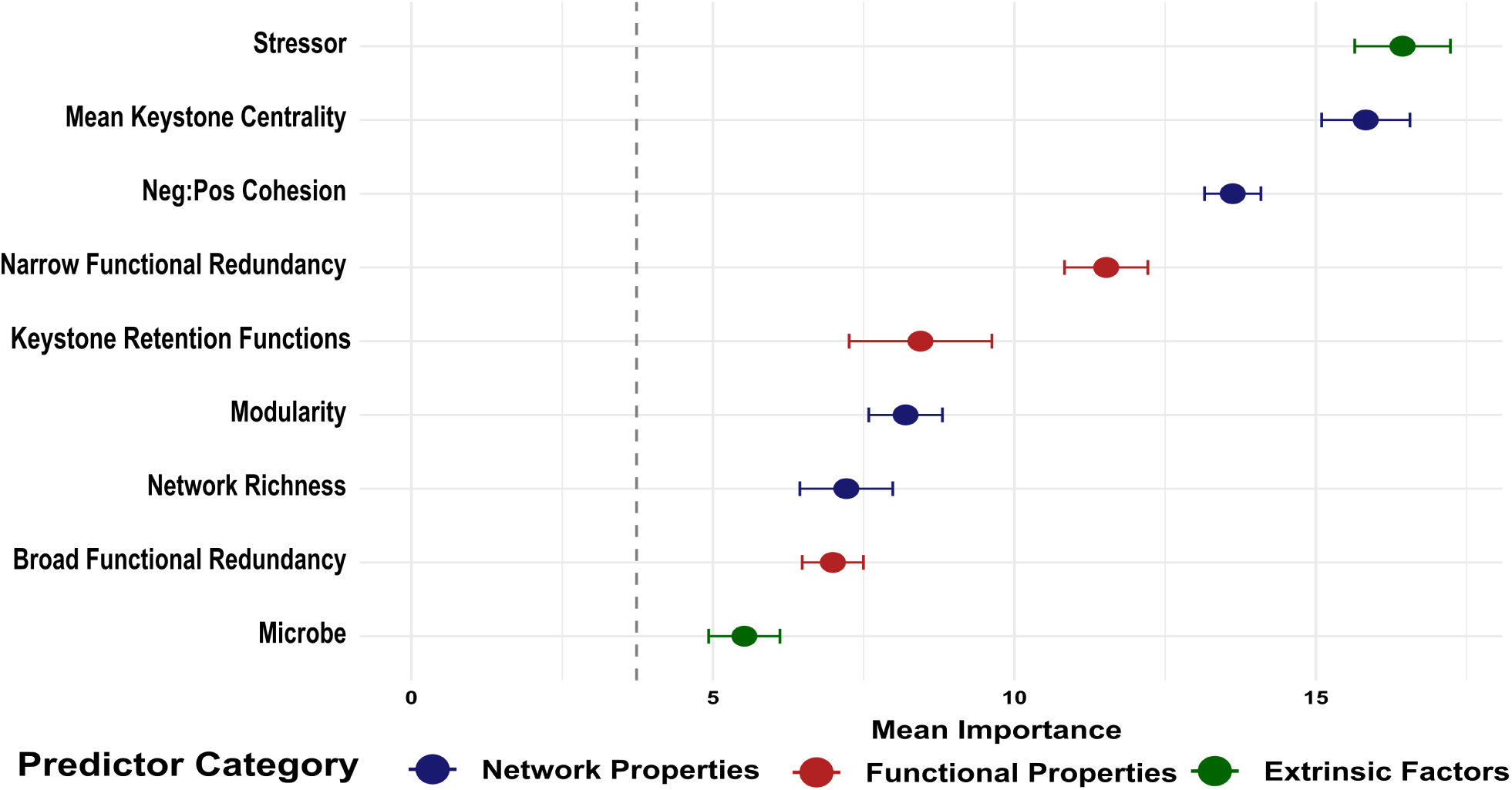
Random forest variable importance for predicting microbial community stability. Mean importance scores from Boruta-confirmed random forest models predicting overall community stability across all three stability dimensions (i.e. structural, functional, compositional). Error bars indicate 95% CI across cross-validation iterations. Points are colored by predictor category: network properties (blue), functional properties (red), and extrinsic factors (i.e. stressor and microbe type) (green). The gray dashed line demarcates the importance threshold below which variables were rejected by Boruta as non-informative (i.e. shadow variables).

For structural shift, more negative interactions (neg:pos cohesion; t = -8.04, p < 0.001, β = - 0.48), higher functional redundancy (broad: t = -4.02, p < 0.001, β = -0.40; narrow: t = -3.93, p < 0.001, β = -0.20), and greater compartmentalization (modularity: t = -2.92, p = 0.004, β = -0.17) were all associated with reduced structural shift (Fig. 3a; Fig. 4a-e). Prokaryotic communities also exhibited greater structural robustness than fungal communities (t = -3.02, p = 0.003, β = - 0.37; Fig. 4e). Together these predictors explained 23% of fixed effects variance and 60% of total model variance including the random effect of study (R²m = 0.23, R²c = 0.60; Supplementary Table 3a). Greater compartmentalization may contain structural disruption within discrete modules, preventing community-wide reorganization under stress. Similarly, a dominance of negative over positive interactions reduces the risk of cascading collapse among taxa that rely upon one another.

For functional shift, higher broad functional redundancy was associated with reduced functional shift (t = -2.94, p = 0.004, β = -0.08; Fig. 3b; Fig. 4f; Supplementary Table 3b), consistent with its protective role in structural shift. In contrast to structural shift, prokaryotic communities exhibited greater functional shift than fungal communities (t = 4.64, p < 0.001, β = 0.08; Fig. 4g), suggesting that the microbiome properties conferring structural robustness do not universally confer functional robustness. Together these predictors explained 20% of fixed effects variance and 44% of total model variance including the random effect of study (R²m = 0.20, R²c = 0.44; Supplementary Table 3b). The consistent protective role of functional overlap across both structural and functional dimensions suggests that maintaining functional overlap within ecological categories is a generalizable strategy for microbiome resilience under climate stress.

Compositional shift was predicted primarily by salinity stressor type (t = 2.64, p = 0.009, β = 0.58; Fig. 3c; Fig. 4h; Supplementary Table 3c), with salinity-stressed communities exhibiting greater compositional turnover than drought-stressed communities (Fig. 4h; Extended Data Fig. 4). The variance explained by the fixed effects was low (R²m = 0.05, R²c = 0.72; Supplementary Table 3c), suggesting that compositional shift is more stochastic and driven by site-specific environmental or historical factors than structural or functional shift. Taken together, our findings reveal that structural and functional microbiome stability are governed by measurable and predictable baseline community properties while compositional turnover remains largely stochastic and context dependent. Critically, microbial communities may maintain their functional capacity under climate stress even as their taxonomic membership changes, a finding with important implications for ecosystem function under future climate scenarios.

## Discussion

Microbiomes underpin the productivity, immunity, and resilience of nearly every plant, animal, and environmental system on Earth, yet no generalizable framework has existed to forecast their stability under climate stress before disturbance occurs. Analyzing over 440 matched network pairs across ∼29,000 microbiomes worldwide, spanning three of the most pressing climate stressors, drought, warming, and salinity, we show that (1) keystone taxa with high functional redundancy, specific metabolic flexibility functions, and central network positions are disproportionately likely to survive stress; (2) at the community level, baseline network and functional properties protect structural and functional stability even as taxonomic composition shifts stochastically; and (3) these relationships hold across host types, ecosystems, and geography. Because these predictive properties can be derived from standard amplicon sequencing data, a low-cost and widely accessible method already used across ecology, agriculture, and conservation, our findings establish baseline microbiome characterization as a viable, scalable tool for forecasting climate vulnerability before disturbance occurs.

We demonstrated that keystone microbe survival is governed by a combination of network connectivity, functional redundancy, and specific retention-associated functions. Highly-connected taxa interact with a greater number of community members, offering more potential pathways of biotic support and buffering against loss^9,25–26^, a property we previously showed drives keystone status during community assembly^23^ and here revealed also governs persistence under stress. We found that keystone retention-associated functions, spanning nitrogen cycling, redox-based energy acquisition, carbon fixation, and stress-response metabolism, converge on a signature of metabolic flexibility, consistent with evidence that flexible metabolic repertoires help microbial communities withstand fluctuating environmental conditions^27–28^, we are conducting a complementary genome-resolved metagenomic analysis to further test this signature. This adds nuance to theory holding that specialists face higher vulnerability to environmental change than generalists^29–31^: rather than being mutually exclusive, ecological specialization and metabolic flexibility can coexist within the same taxon, and it is specifically the keystones retaining generalist-like flexibility that persist, underpinning broader community-level resilience.

Baseline network and functional properties predicted structural and functional shift with substantial explanatory power, using aggregate metrics of functional diversity and redundancy that are more robust to prediction uncertainty than fine-grained, individual enzyme-level inferences. This is consistent with evidence that microbial community properties, rather than species richness alone, govern the resistance of ecosystem multifunctionality to global change^32^. Compositional shift, in contrast, remained largely unpredictable, driven primarily by stressor identity rather than any measured community property, consistent with the broader recognition that stochastic processes such as drift and dispersal can play a substantial role in shaping microbial community composition^33^, and with prior work showing that nutrient cycling and community composition can decouple under aridity stress at a global scale^34^. This asymmetry extends a body of work showing that taxonomic and functional variation constitute largely independent axes in microbial communities^35^ by demonstrating that this decoupling has a temporal, stress-response dimension as well: communities can maintain predictable structural and functional trajectories under climate stress, plausibly through mechanisms such as compartmentalization limiting the spread of perturbation across the network^36^, even as the specific taxa comprising them turn over stochastically. This finding reframes what stability should mean for a microbiome under climate stress: rather than requiring the persistence of particular taxa, resilience may be better defined by the persistence of the structural and functional roles those taxa fill, roles that our results show are forecastable from baseline properties even when the identity of the organisms filling them is not.

The predictive relationships we identified held consistently across host types, ecosystems, and stressor identities, and stable and unstable networks were distributed globally without geographic clustering. Climate and geography still matter; a given magnitude of warming, drought, or salinization can represent a far more severe departure from baseline conditions in some regions than others, since a few degrees of warming may be mild in an already warm system but push a historically cold system past critical phase-change thresholds, such as permafrost thaw^37^. Our paired design captures this relative shift directly, comparing each stressed network only to its own baseline, rather than relying on an absolute stressor value with different biological meaning across climates. That the same baseline properties predicted stability regardless of location, despite this variation in what a given stressor represents locally, underscores the strength of the framework, distinguishing it from approaches reliant on region-specific climate projections or species distribution models, which are often difficult to transfer between systems^38–39^. A trait-based framework, by contrast, can be broadly applied across microbiomes with baseline amplicon sequencing data, complementing climate-mediated frameworks used in other systems, where the biodiversity-stability relationship itself has been shown to vary with climate^40^. This generalizability elevates baseline microbiome characterization from a descriptive tool to a genuinely predictive one across the diverse ecosystems and stressors we tested. To make this framework immediately usable, we provide our trained predictive models directly, allowing researchers to apply them to their own baseline microbiome data without re-running the full analytical pipeline described here.

While our findings establish that baseline network and functional properties predict microbiome stability across a globally diverse dataset spanning bacteria, archaea, and fungi, future progress will depend on the pace at which the field generates new kinds of empirical data. Multi-stressor experiments, longitudinal sampling across full stress-recovery trajectories, and manipulated synthetic communities with defined network properties remain comparatively rare in the public sequence archives from which datasets like ours are built. As these designs become more common, they will make it possible to test whether the same baseline properties predict resilience under compound climate extremes, over longer successional timescales, and under direct experimental manipulation of network structure, further strengthening and extending the framework established here. Likewise, as sequencing of microeukaryotes and viruses across diverse systems continue to expand, and as genome-resolved metagenomic data become more widely available, these resources will offer additional opportunities to test the generality and mechanistic basis of the predictors we identified. Building on this foundation will sharpen an already powerful tool: because our framework requires only standard, low-cost amplicon sequencing collected before disturbance occurs, it is immediately deployable across the ecosystems, agricultural systems, and conservation priorities where microbiome resilience matters most, offering a practical path toward safeguarding the microbial foundations of life on Earth as climate change accelerates.

## Methods

### Microbiome data collection

Datasets of control and stressed microbiomes were obtained from the Sequence Read Archive within the National Center for Biotechnology and Information (NCBI) which contains publicly available microbiome sequencing raw reads and their associated metadata (accessed 28 June 2023). We utilized NCBI’s search engine Entrez to search for datasets by using pairwise combinations of a microbial search term “community, microbiome, prokaryotic, fungal, eukaryotic, bacterial, 16S, ITS, 18S” and a stressor search term “drought, salinity, warming.” We focused on drought, warming, and salinity because they are among the most pervasive and consequential climate stressors driving biodiversity loss and ecosystem destabilization worldwide, making them a priority for developing generalizable predictors of microbiome resilience^6^. We used the following filtering criteria to narrow down the identified datasets and to maintain comparability among studies: 1) the sequencing platform used was Illumina, 2) datasets had a minimum of 8 replicates each for control and stressed groups, and 3) metadata included the stressor type, association-type (i.e. free-living, host-associated), and geographic location of study (i.e. latitude, longitude). This initial search was supplemented with more recent studies meeting the same inclusion criteria identified after the original search date. When required metadata were not available in the associated publication or NCBI record, we contacted the corresponding authors directly to obtain the missing information.

This filtering followed standard practices in microbiome meta-analysis which allowed us to compare sequences of like types across diverse studies to reduce biases^41–43^ and enabled us to apply the same quality control and bioinformatics pipelines on all raw reads obtained so that the generated ASVs were standardized for all analyses^43–45^. Rather than restricting our scope to a single host, habitat, or region, we deliberately compiled microbiomes from host-associated and free-living systems across terrestrial and marine environments worldwide, capturing the full breadth of ecological contexts in which climate stress acts on microbial communities. Realizing generalizable predictors from such heterogeneous systems required a design that could isolate the signal of stress exposure itself; therefore, only studies with a paired control versus stressed design, in which both groups were sampled from the same system and differed only in stress exposure were retained. Each stressed network was thus compared only to its own baseline counterpart, allowing global variation in host, habitat, and geography to serve as a test of generalizability rather than a source of noise. Our final dataset consisted of 110 studies spanning drought, warming, and salinity stressors ranging from alpine permafrost^46^ to subtropical agroecosystems^47^ to coral reefs^48^. A list of the NCBI project IDs and corresponding publication references for all datasets used is available in Supplementary Table 1.

Amplicon sequencing was selected as the primary basis for this analysis for three reasons specific to the aims of this study. First, robust co-occurrence network construction requires a minimum of 8 replicates per control and stressed group^49^; datasets meeting this replication threshold together with a paired control-versus-stressed design were substantially more available among amplicon studies than among shotgun metagenomic studies with genome-resolved data (metagenome-assembled genomes; MAGs), reflecting the comparatively lower cost and broader adoption of amplicon sequencing across ecological and environmental microbiology^50^. This disparity in available replication is illustrated directly by our genome-resolved comparison: despite an extensive literature search, only 10 studies with paired MAG data met minimal inclusion criteria, compared to 110 amplicon-based studies, and MAG sample sizes among most of these were insufficient to support network-level analysis at all (Supplementary Table 1). Second, the amplicon-based approach allowed inclusion of both prokaryotic and fungal communities (16S, ITS, and 18S), whereas MAG recovery from shotgun metagenomic data is overwhelmingly restricted to prokaryotic taxa^51^. Therefore, genome-resolved data could not replicate the cross-kingdom comparison central to testing the generalizability of our framework. Third, the breadth of amplicon datasets available across host types, habitats, and geographic regions was essential to testing whether network structural and functional predictors of stability generalize across ecological contexts, rather than being an artifact of any single system, a scope not achievable with the far more limited pool of MAG-based studies. To directly assess whether amplicon-derived functional patterns are consistent with genome-resolved data, we additionally conducted a targeted complementary analysis using metagenome-assembled genomes (MAGs) from studies with available shotgun metagenomic sequencing (see section *Genome-resolved assessment of functional patterns*, below).

### Quality control and bioinformatics pipeline

Raw 16S, ITS, and 18S nucleotide sequence data and corresponding metadata were downloaded from NCBI using q2-fondue^52^. Sequences were processed following the standardized bioinformatics pipeline previously described^53^. In brief, FIGARO^43^ was used to determine optimal quality trim parameters, the QIIME2 pipeline^54^ was used to remove low-quality bases and join paired-end reads, and the DADA2 algorithm^55^ was used to generate amplicon sequence variants (ASVs). ASVs were taxonomically classified using pretrained Naïve Bayes classifiers from the SILVA138.2 V4 region database for prokaryotes^56^ and the UNITE v.10.0 ITS region database for fungi^57^. Samples with fewer than 2,000 reads and ASVs with relative abundances below 0.25% were discarded to remove samples with insufficient sequencing depth and to avoid inflating diversity estimates with contaminant or spurious sequences^18,58^. Because some studies contributed multiple control-stress pairs originating from distinct sample sources, hosts, or habitats, ASV count data were separated by study, sample source, microbial kingdom, and treatment (control versus stressed) for all downstream analyses. Stochastic ASVs, defined as those present in fewer than 5% of samples^59–60^ (or found in only a single sample when sample size was < 20), were removed from the count data prior to network construction, as stochastic taxa have been shown to have little influence on microbial community structure^23^.

### Microbiome network construction

The minimum sample size requirement of 8 biological replicates for each control and stressed group, applied during dataset selection, also ensured that edge correlations within each network could be estimated reliably^49^. Each control and stressed dataset was used to construct a pair of co-occurrence networks using FastSpar^61^, a parallelized implementation of SparCC that accounts for the compositional sparsity characteristic of microbiome sequencing data^62^, run with default parameters. We selected SparCC over alternative network inference methods because it maintains comparatively low false discovery rates at sample sizes under 1,000 and, unlike several other approaches, does not impose constraints that could obscure highly connected hub taxa, a structural feature central to our analysis of keystone retention^63–64^. SparCC is also widely adopted for microbiome network reconstruction across plant, soil, and other environmental systems^65^ and has been validated as an accurate approach for capturing network structure in natural communities^18,23,66^. This approach yielded 441 paired control and stressed networks (882 networks total) across the 110 studies retained for analysis.

### Calculation of network structural predictors

For each of the 882 networks, we calculated modularity, negative:positive cohesion, a composite centrality index, and network richness. Modularity, which reflects the proportion of connections occurring within versus between network modules, was calculated using the Clauset–Newman–Moore greedy modularity detection algorithm^67^. In biological networks, modularity values typically range from 0 to 1, with higher values indicating greater compartmentalization of the network into discrete interaction modules^16^. Cohesion was calculated by weighting the correlation strength of each taxon’s edges by its relative abundance and normalized by its number of edges, then summed separately across positive and negative edges^17^. The ratio of summed negative to summed positive cohesion was used as the metric for each network. The absolute value of this ratio was used in subsequent analyses, with values greater than 1 indicating that negative interactions contributed more strongly to network cohesion than positive interactions, values less than 1 indicating the reverse, and a value of 1 indicating an equal contribution of negative and positive interactions. A composite centrality index was calculated, and taxa were binned into central and intermediate taxa as previously described^23^; briefly, degree, betweenness, closeness, and eigenvector centrality were combined into a single centrality index using principal components analysis, and the top 5% of taxa by centrality index were classified as central. Network richness was calculated as the number of taxon nodes present in the network.

### Keystone taxon identification and retention

Keystone taxa were defined as those classified in the central centrality tier^23^ (for more information refer to *Calculation of network structural predictors* above). For each control-stressed network pair, keystone taxa identified in the control network were used to define a baseline keystone pool, and a keystone taxon was considered retained if it was still present in the paired stressed network. We used presence-absence rather than abundance-based comparisons because local extinction, rather than a change in relative abundance, is the outcome most directly relevant to recovery^68–69^: a keystone taxon that persists at reduced abundance retains the potential to re-establish its interactions, whereas one that is lost entirely does not. Keystone persistence was calculated at the level of the individual taxon (present or absent in the stressed sample) and aggregated to the network level as the percentage of baseline keystone taxa retained following stress, calculated as (1 − Jaccard dissimilarity between the baseline keystone pool and its paired stressed sample) × 100%.

### Calculation of network functional predictors

Functional potential was predicted for both prokaryotic and fungal amplicon data using PICRUSt2, which infers community-level gene family abundances by placing sequences onto a reference phylogeny and mapping to characterized reference genomes^70^. Each ASV was required to align to at least 50% of its length to the reference prior to phylogenetic placement, following PICRUSt2 recommendations for analyses that include fungal communities^71^. Predicted gene families were mapped to enzyme commission (EC) numbers using the expanded PICRUSt2-SC database^71^, rather than KEGG Orthology, because EC classifications provide more consistent coverage across fungal reference genomes while retaining good coverage for bacterial genomes, allowing functional potential to be compared directly across kingdoms^70,72^. A substantial fraction of microbial functional diversity remains uncharacterized in reference databases, which can limit the reliability of functional predictions in some microbiome studies^70^. This was not a major limitation for our dataset, however, as an average of 84% of ASVs across individual sequencing samples, prior to network construction, were successfully matched to the PICRUSt2-SC database.

For each sample, EC hits were combined across all detected taxa to generate a sample-level presence/absence profile of functional potential. Rather than restricting this profile to a subset of common functions, every EC number predicted for a given sample was retained, ensuring that rare and less common functions, which can be biologically important but are often excluded by prevalence filtering^73^, were not lost from downstream analyses. EC functions present in at least 75% of samples were classified as broad, and all remaining functions were classified as narrow; functional diversity (the number of unique EC functions present) and functional redundancy (the number of EC hits divided by functional diversity) were calculated separately for the overall, broad, and narrow EC sets.

To identify EC functions associated with keystone retention, we used indicator species analysis^74^ to relate individual EC functions carried by keystone taxa to whether those taxa were retained or lost following stress. EC functions were classified as significantly associated with retention if they had an adjusted p-value ≤ 0.05 and an indicator statistic ≥ 0.15, a threshold selected to balance retention of biologically meaningful functions against the risk of spurious associations given the large number of EC functions tested. This retention-associated EC set was used to calculate a per-keystone retention EC proportion (the proportion of a keystone taxon’s EC functions belonging to the retention-associated set) and a network-level retention EC score (for each retention-associated EC function, the proportion of a network’s baseline keystones carrying that function, averaged across the full set of retention-associated EC functions), both of which were used as predictors in subsequent models.

### Genome-resolved assessment of functional patterns

Because the 10 MAG studies included in this analysis originally employed differing bioinformatic pipelines for MAG abundance estimation and functional annotation, we re-derived both directly from each study’s raw sequencing data using a single, standardized pipeline, ensuring that MAG abundance and functional annotation were calculated identically across all studies rather than inheriting each study’s original, non-uniform methodology. MAG abundance was derived by mapping each study’s raw reads to their corresponding MAGs using CoverM^75^. Functional annotation was performed de novo for all MAGs using DRAM^76^; KEGG Orthology (KO) annotations were converted to enzyme commission (EC) numbers using the KEGG ko2ec reference table^77^ to enable direct comparison with amplicon-derived PICRUSt2-SC EC predictions. MAG abundance was used to weight EC annotations, producing an abundance-weighted EC-by-sample matrix. To identify EC functions associated with control versus stressed samples, we used indicator species analysis, relating individual EC functions to sample condition using the same approach applied to the amplicon-derived indicator analysis (see *Keystone taxon identification and retention*, above). Because the limited sample sizes available for MAG-based studies (see above) precluded a keystone-retention framework analogous to the amplicon-based analysis, this indicator analysis instead used condition (control versus stressed) as the grouping variable, representing the closest available parallel to the amplicon-derived approach. For functional redundancy, this same EC-by-sample matrix was converted to presence/absence, and redundancy was calculated using the same approach applied to amplicon-derived EC data (see *Calculation of network functional predictors*, above), enabling direct comparison against the amplicon-derived functional redundancy metric.

### Calculation of network shift and stability

To quantify the overall magnitude of structural change induced by stress, we calculated a dissimilarity index for modularity, cohesion, and network richness as the absolute difference between the stressed and control network value, normalized by their sum. Structural shift was calculated as the mean of these three property-level dissimilarity indices, expressed as a percentage. Functional shift was calculated as the Jaccard dissimilarity between the EC presence/absence profile of the control sample and its paired stressed sample, referenced against the baseline (control) EC set, expressed as a percentage. Compositional shift was calculated using the same baseline-referenced Jaccard approach, applied to the full taxonomic composition of central and intermediate taxa (i.e. all taxa retained within network construction thresholds) rather than to EC functions or keystones specifically, expressed as a percentage. An overall shift score was calculated for each network pair as the mean of structural, functional, and compositional shift. Network pairs with an overall shift score below 20% were classified as stable, and all others were classified as not stable.

### Statistical analyses

To test whether individual keystone attributes predicted keystone persistence, we fit a binomial generalized linear model with keystone retention (retained versus lost) as the response variable and keystone EC redundancy and centrality as predictors. We additionally tested for differences in centrality and EC redundancy between retained and lost keystones using linear mixed-effects models, with persistence status as a fixed effect and study included as a random intercept to account for non-independence among keystones drawn from the same study. To evaluate whether these keystone-level attributes, aggregated to the network level, predicted the overall proportion of keystones retained within a network, we fit a linear model with the percentage of baseline keystones retained as the response variable and mean keystone centrality, the network-level retention EC score, and mean keystone EC redundancy as predictors.

To further evaluate how well keystone attributes predicted individual keystone persistence, we constructed a random forest classification model^78^ using keystone status (retained versus lost) as the response variable, with keystone EC redundancy, keystone centrality, the per-keystone retention EC proportion, stressor type, and microbial kingdom as candidate predictors. Prior to model fitting, Boruta feature selection^79^ was used to identify predictors significantly more informative than randomly permuted shadow predictors; any predictors that could not be conclusively classified as confirmed or rejected within the algorithm’s iterations were resolved by comparing their importance to that of confirmed predictors. Model performance was evaluated using 10-fold cross-validation^80^, with the area under the receiver operating characteristic curve (AUC) as the evaluation metric; AUC values of 0.7+ are considered acceptable discrimination^81^.

Linear mixed-effects models (LMMs) were used to test whether baseline network structural and functional properties predicted the magnitude of structural, functional, and compositional shift following stress. Each model included modularity, absolute value neg:pos cohesion, network richness, overall functional redundancy, mean keystone centrality, the network-level retention EC score, and mean keystone EC redundancy as fixed effects, along with stressor type and microbial kingdom as fixed effects to account for baseline differences among stressors and taxonomic groups. Study was included as a random intercept to account for non-independence among network pairs drawn from the same study. Three separate models were fit, one for each shift metric as the response variable; the functional shift model used a log-transformed response to meet model assumptions of normality. Variance inflation factors were checked to assess multicollinearity among predictors, residual normality was assessed using the Shapiro-Wilk test, and marginal and conditional R² were calculated for each model. To evaluate predictors of network stability following stress, we constructed a random forest classification model using the same approach described above, with stability category (stable versus not stable, defined above) as the response variable and the same set of structural, functional, and keystone-level predictors used in the LMMs as candidate predictors. AUC values for this model also fell within the acceptable range.

To make these models directly reusable, the trained community stability and keystone persistence random forest models were exported as .rds objects and will be released upon publication. Applying these models to new baseline microbiome data requires calculating the same set of network structural, functional, and keystone-level predictors described above from the new dataset; a description of the required input format is provided alongside the model files as a README.

## Author Contributions

A.H.R., A.L.R., V.W.L., J.C.F., E.M.H., D.J.H., and M.E.A. conceptualized the study and contributed to methodology development. A.H.R. and D.J.H. developed the bioinformatics pipelines used in this study. A.H.R. performed all formal analyses, constructed all figures, and wrote the original draft. All authors contributed to data validation, investigation, and curation. A.H.R. and M.E.A. administered the project. M.E.A. supervised the research and acquired funding. All authors reviewed and edited the manuscript.

## Data and Code Availability

All raw sequence data used in this study are publicly available through the NCBI Sequence Read Archive. Project IDs for all datasets are listed in Supplementary Table 1. Supplementary files are available with the published article. Our trained predictive models, including the random forest models for community stability and keystone persistence as described in Methods, are available as .rds objects upon publication, along with a description of the required input data format for their application to new datasets as a README. All additional processed data and analysis code are available upon publication.

## Acknowledgements

We are grateful to members of the Afkhami and Searcy labs and A. Igwe for valuable discussions and feedback throughout this project. Computational analyses were performed on the Pegasus high-performance computing cluster at the University of Miami.

