## Supplementary_Figures_and_Tables for "A global signature of microbiome resilience to climate stress"

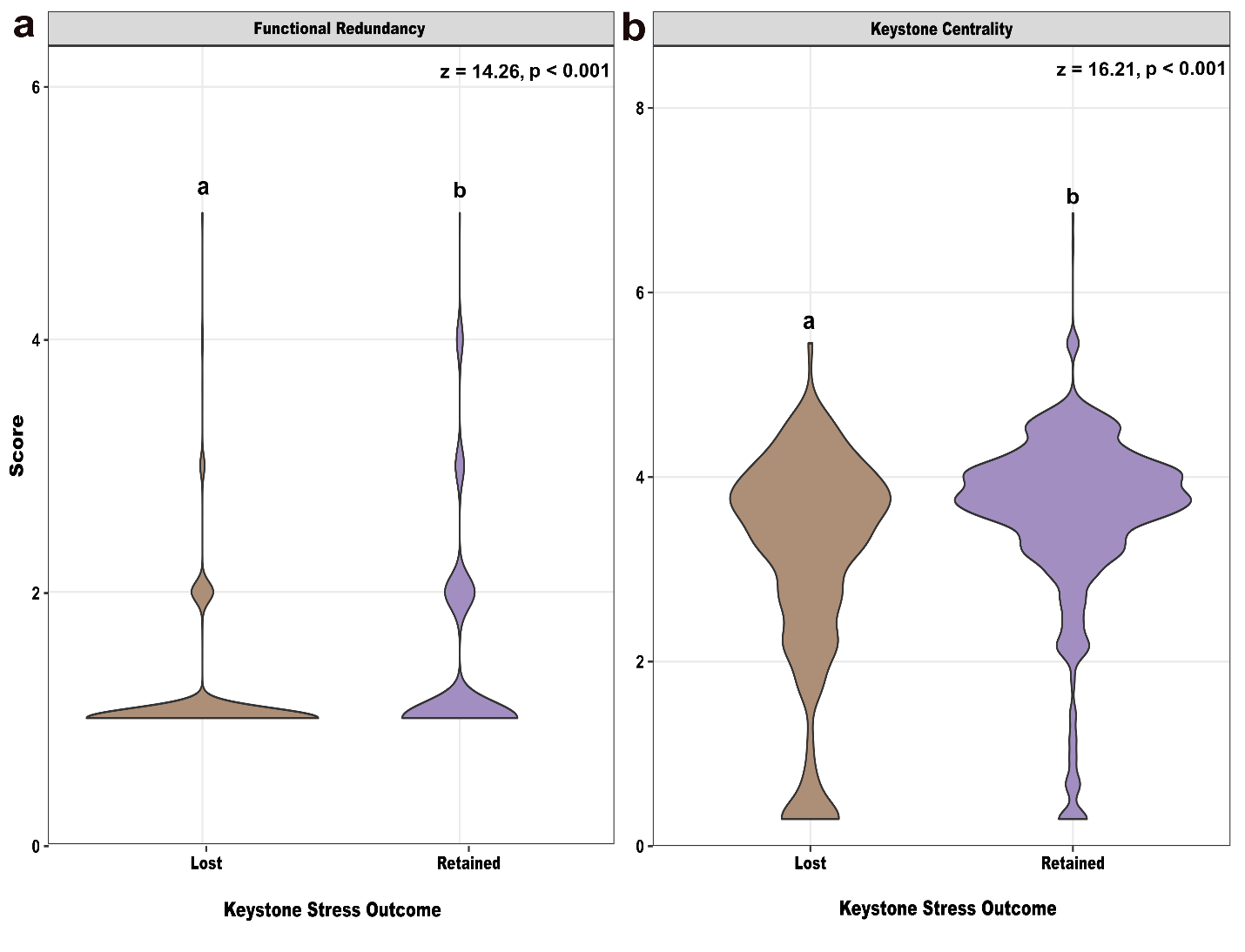

**Extended Data Fig. 1: Retained keystone taxa exhibit higher functional redundancy and centrality than lost keystones.** Raw score distributions of **(a)** functional redundancy and **(b)** keystone centrality for keystone taxa lost or retained following climate stress events. Statistics shown are z-values and p-values from logistic regression model.

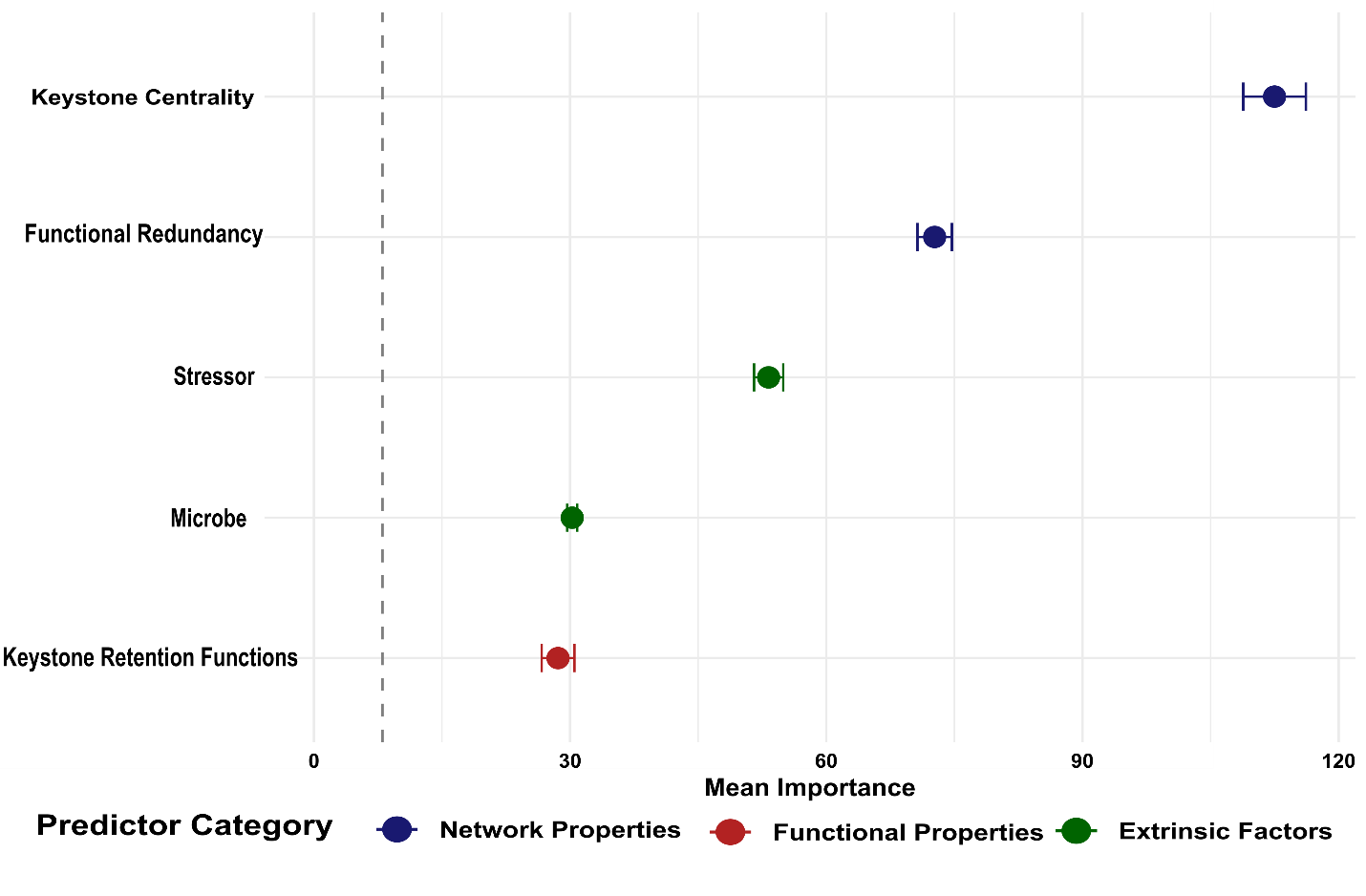
**Extended Data Fig. 2: Random forest variable importance for predicting keystone microbe persistence during stress.** Mean importance scores from Boruta-confirmed random forest models predicting individual keystone persistence during stress. Error bars indicate 95% CI across cross-validation iterations. Points are colored by predictor category: network properties (blue), functional properties (red), and extrinsic factors (i.e. stressor and microbe type) (green). The gray dashed line demarcates the importance threshold below which variables were rejected by Boruta as non-informative (i.e. shadow variables).

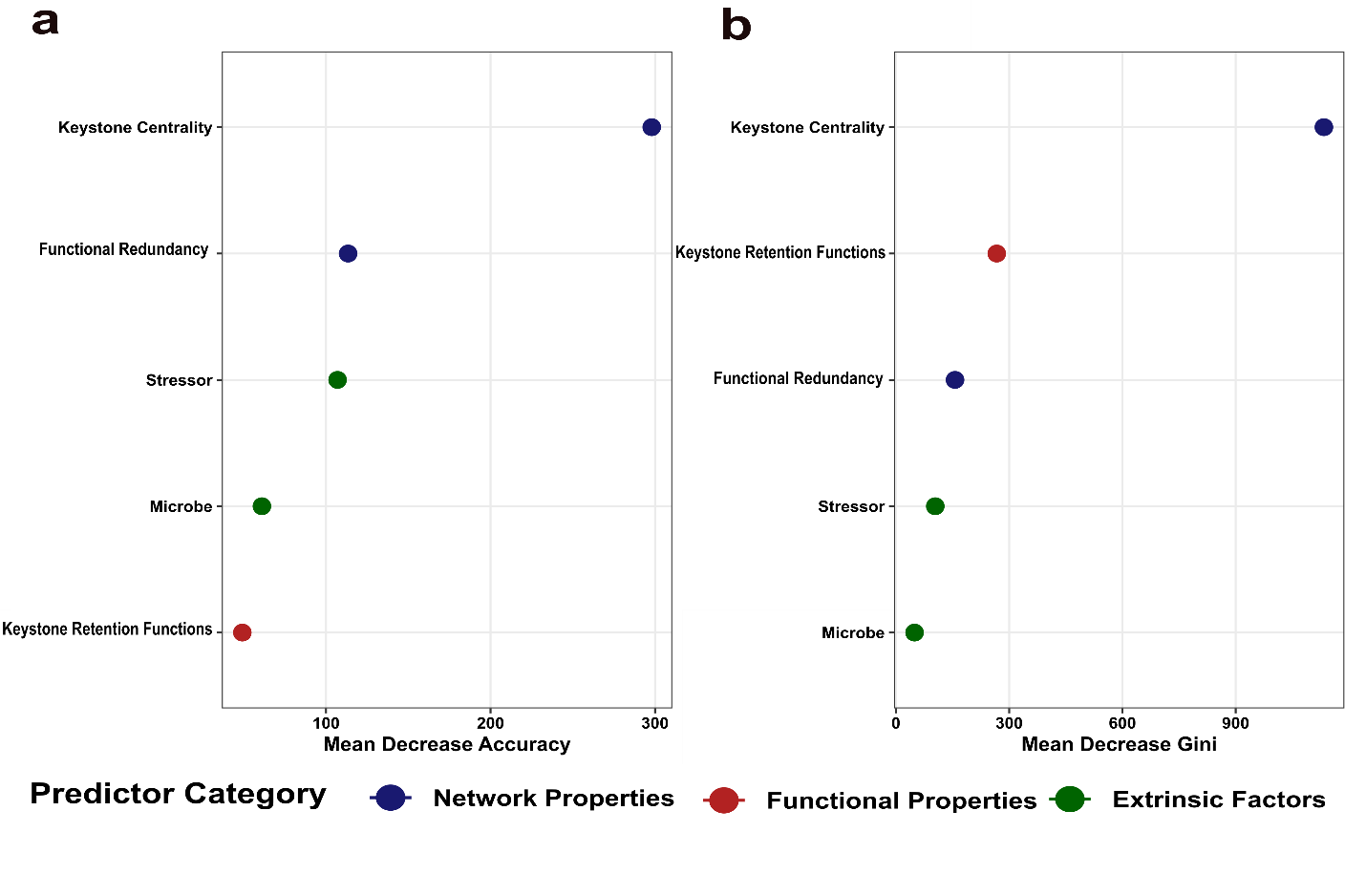
**Extended Data Fig. 3: Random forest variable importance metrics for keystone persistence model.** **(a)** Mean decrease accuracy and **(b)** mean decrease gini for predictors in the keystone persistence random forest model. Points are colored by predictor category: network properties (blue), functional properties (red), and extrinsic factors (green). Concordance between metrics indicates robust variable importance independent of the measurement approach.

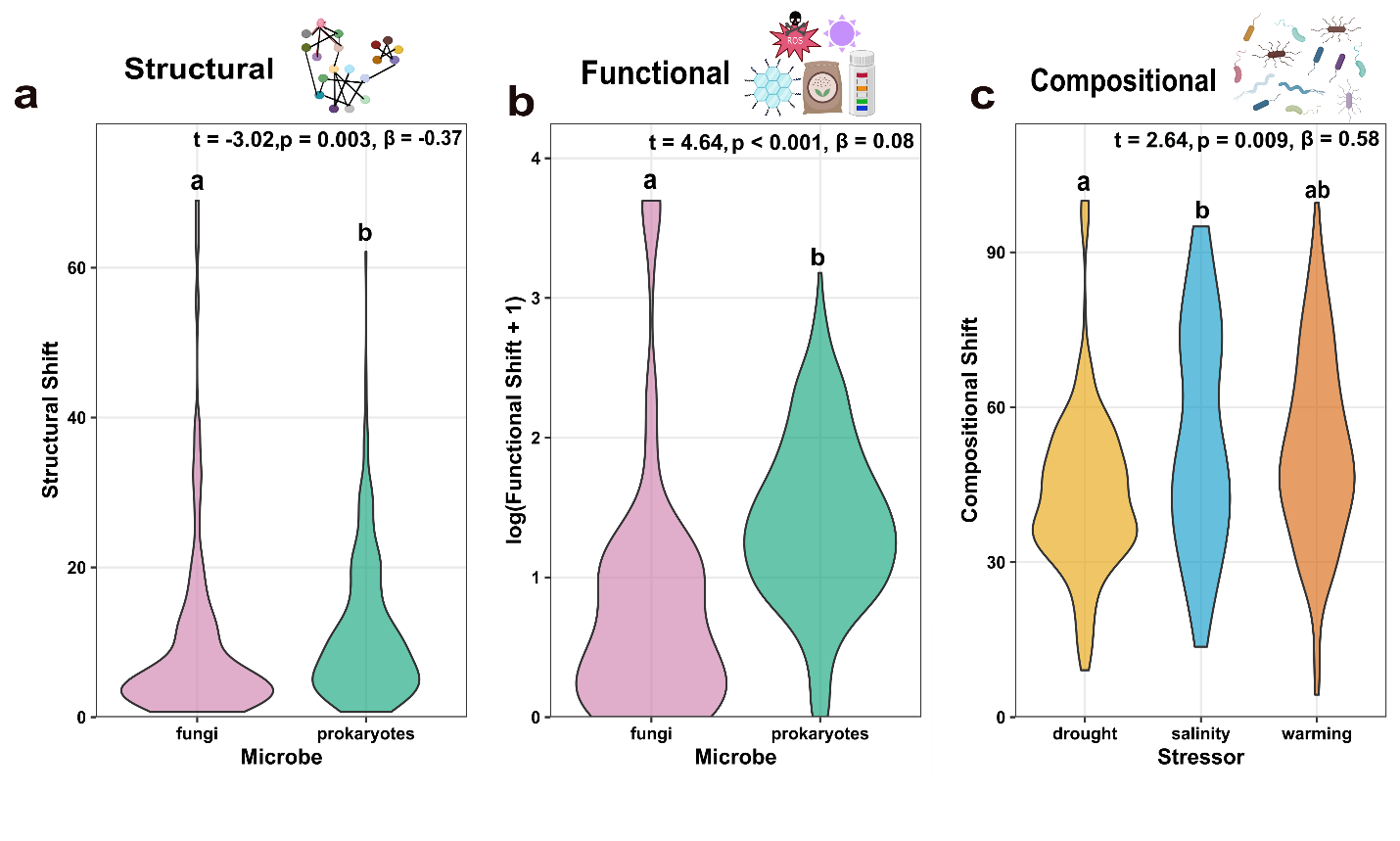

**Extended Data Fig. 4: Raw distributions of community shift by microbe and stressor type.** Violin plots showing the full distribution of **(a)** structural, **(b)** functional, and **(c)** compositional shift after stress. Statistics shown are t-values, p-values, and standardized beta coefficients from linear mixed models.

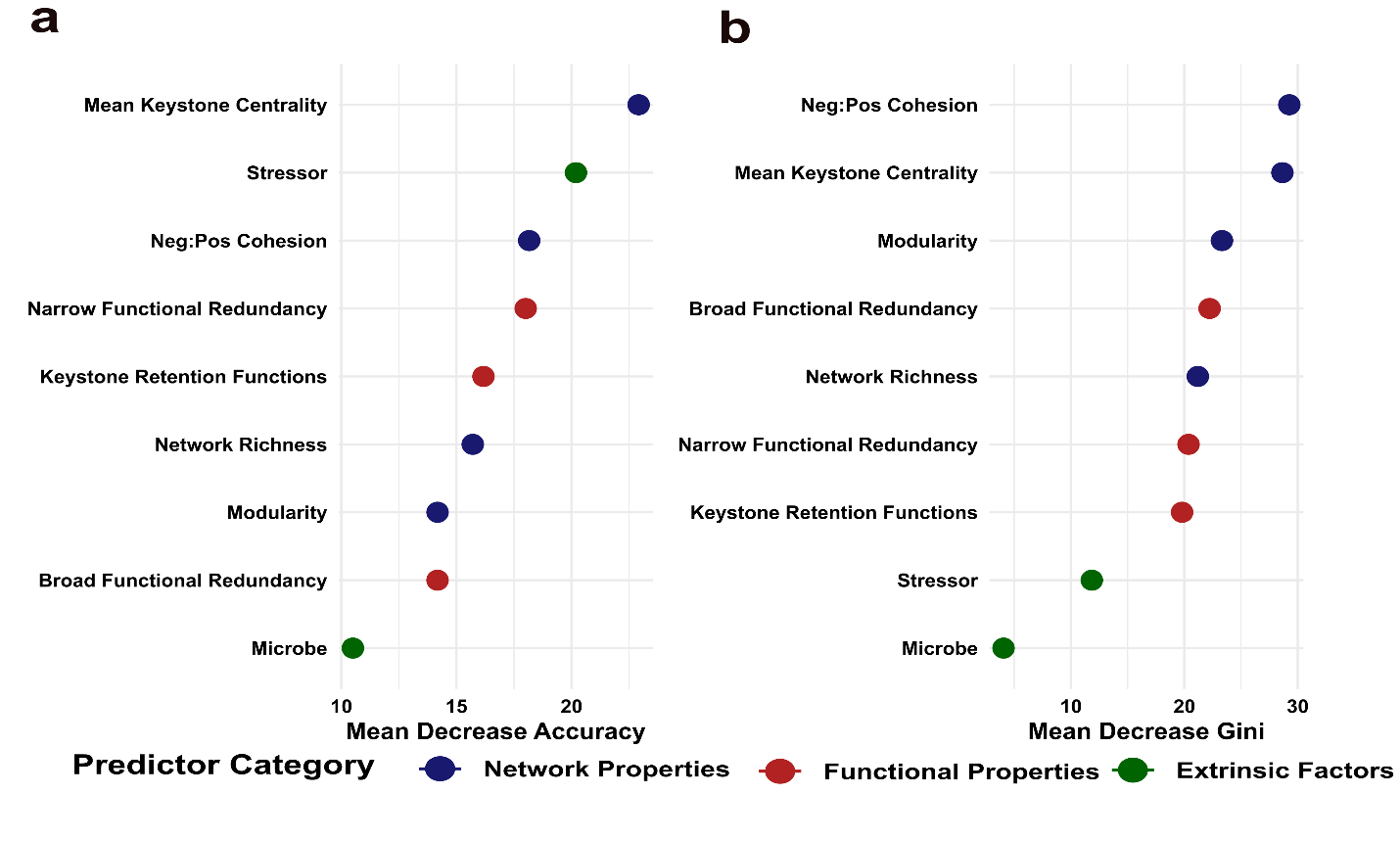

**Extended Data Fig. 5: Random forest variable importance metrics for community stability model. (a)** Mean decrease accuracy and **(b)** mean decrease gini for predictors in the community stability random forest model. Points are colored by predictor category: network properties (blue), functional properties (red), and extrinsic factors (green). Concordance between metrics indicates robust variable importance independent of the measurement approach.

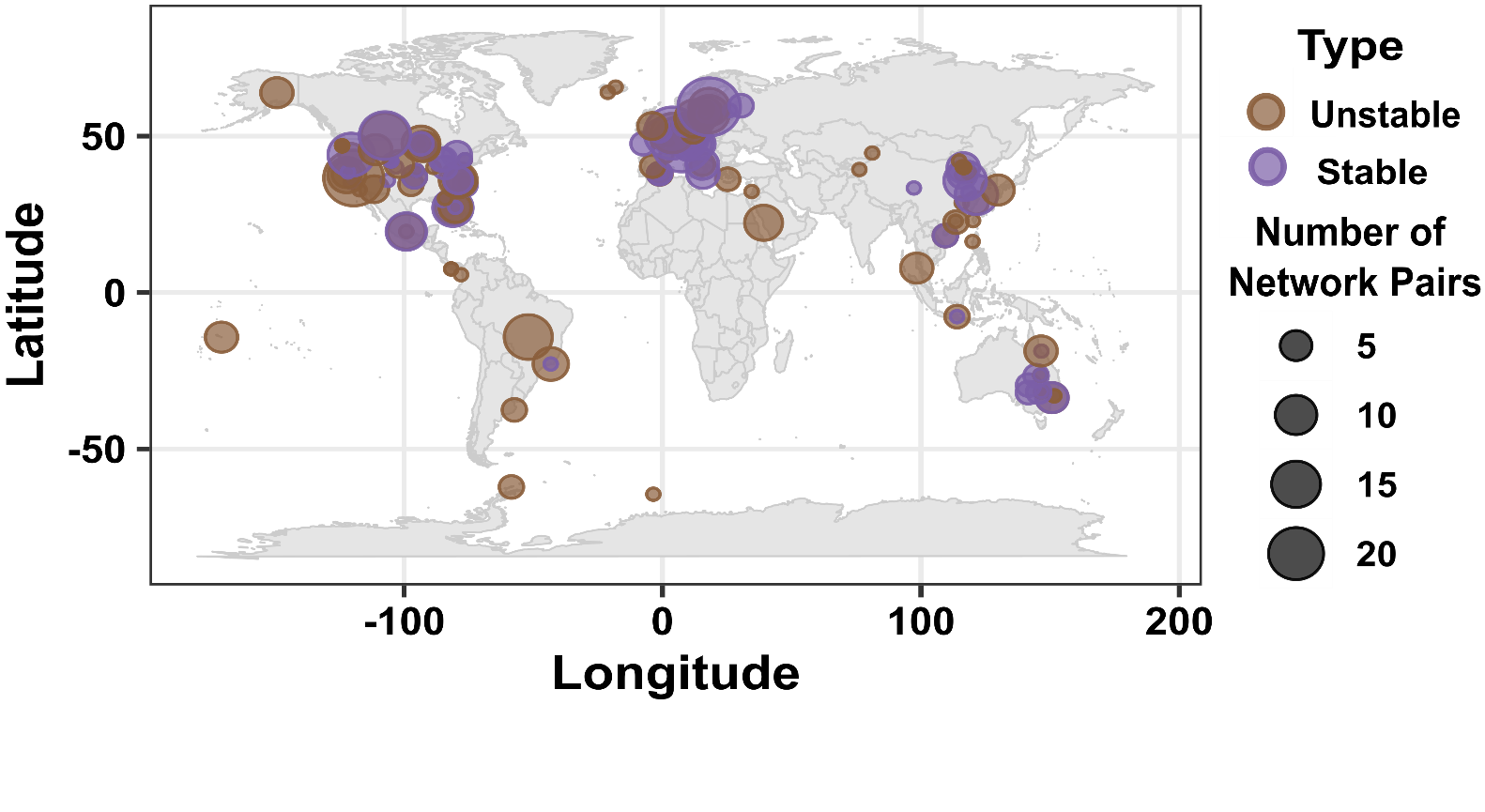
**Extended Data Fig. 6: Stable and unstable microbial communities are distributed globally, suggesting microbiome community properties rather than geography predict stability under climate stress.** Geographic distribution of network pairs classified as stable (n = 228; purple) or unstable (n = 213; brown) following climate stress events, based on the community stability threshold derived from the random forest model. Circle size indicates the number of network pairs per location.

**Supplementary Tables:**

**Supplementary Table 1: Data sources summary (see Supplementary_Table_1.xlsx).** For each of the 110 studies included across three climate stressors the geographic location (country), association type (free-living or host-associated), sample type, microbe type (prokaryotes, fungi, or both), number of microbiome samples, number of network pairs, and NCBI BioProject accession number is provided. In addition, all citations and DOIs are provided for all data sources utilized. Studies are highlighted in yellow, blue, and red to correspond to drought, salinity, and warming stressors, respectively.

**Supplementary Table 2: Keystone taxa model summary. (a)** Logistic regression model predicting individual keystone persistence (lost vs. retained) during stress events. **(b)** Linear model predicting the percentage of keystones retained per network, with standardized beta coefficients and 95% confidence intervals. **(c)** Random forest model performance metrics from 10-fold cross-validation predicting keystone persistence. Significance codes: *** p < 0.001, ** p < 0.01, * p < 0.05.

| ***(a) Logistic Regression*** | | | | | |
| --- | --- | --- | --- | --- | --- |
| **Predictor** | **Estimate** | **SE** | **z** | **p** | **Sig.** |
| Intercept | -2.043 | 0.099 | -20.55 | <0.001 | *** |
| Functional Redundancy | 0.693 | 0.049 | 14.26 | <0.001 | *** |
| Keystone Centrality | 0.399 | 0.025 | 16.21 | <0.001 | *** |

| ***(b) Keystones Retained (%) Linear Model*** | | | | | | | |
| --- | --- | --- | --- | --- | --- | --- | --- |
| **Predictor** | **Estimate** | **SE** | **t** | **p** | **Sig.** | **Std. β** | **95% CI** |
| Mean Keystone Centrality | 4.103 | 1.696 | 2.42 | 0.016 | * | 0.12 | [0.02, 0.22] |
| Mean Proportion of Keystone Retention Functions | 99.830 | 28.121 | 3.55 | <0.001 | *** | 0.17 | [0.08, 0.27] |
| Mean Functional Redundancy | 6.663 | 3.224 | 2.07 | 0.039 | * | 0.10 | [0.01, 0.20] |

| ***(c) Random Forest (10-fold CV)*** | |
| --- | --- |
| **Metric** | **Value** |
| AUC | 0.775 |
| Accuracy | 0.713 |
| Optimal mtry | 6 |

**Supplementary Table 3. Community stability model summary.** Linear mixed models predicting **(a)** structural, **(b)** functional, and **(c)** compositional shifts after climate stress, with study as a random effect. Significance codes: *** p < 0.001, ** p < 0.01, * p < 0.05. Marginal R² (R²m) reflects variance explained by fixed effects; conditional R² (R²c) reflects total variance explained including random effects. **(d)** Random forest model performance from 10-fold cross-validation predicting community stability (stable vs. not stable).

| ***(a) Structural Shift Linear Mixed Model*** | | | | | |
| --- | --- | --- | --- | --- | --- |
| **Predictor** | **Estimate** | **SE** | **t** | **p** | **Sig.** |
| Modularity | -12.589 | 4.315 | -2.918 | 0.004 | ** |
| Neg:Pos Cohesion | -24.500 | 3.048 | -8.037 | <0.001 | *** |
| Network Richness | 0.001 | 0.005 | 0.298 | 0.766 |  |
| Broad Functional Redundancy | -0.058 | 0.015 | -4.019 | <0.001 | *** |
| Mean Keystone Centrality | 0.394 | 0.618 | 0.637 | 0.525 |  |
| Keystone Retention Functions | -12.839 | 8.782 | -1.462 | 0.145 |  |
| Narrow Functional Redundancy | -4.740 | 1.206 | -3.930 | <0.001 | *** |
| Stressor (salinity) | 1.062 | 1.912 | 0.555 | 0.580 |  |
| Stressor (warming) | 0.095 | 1.336 | 0.071 | 0.944 |  |
| Microbe (prokaryotes) | -3.752 | 1.241 | -3.023 | 0.003 | ** |
| *Marginal R² = 0.232, Conditional R² = 0.602* | | | | | |

| ***(b) Functional Shift Linear Mixed Model*** | | | | | |
| --- | --- | --- | --- | --- | --- |
| **Predictor** | **Estimate** | **SE** | **t** | **p** | **Sig.** |
| Modularity | 0.213 | 0.344 | 0.618 | 0.537 |  |
| Neg:Pos Cohesion | -0.127 | 0.242 | -0.526 | 0.599 |  |
| Network Richness | 0.001 | 0.000 | 1.641 | 0.102 |  |
| Broad Functional Redundancy | -0.003 | 0.001 | -2.938 | 0.004 | ** |
| Mean Keystone Centrality | -0.033 | 0.050 | -0.653 | 0.514 |  |
| Keystone Retention Functions | -0.737 | 0.720 | -1.024 | 0.307 |  |
| Narrow Functional Redundancy | 0.098 | 0.098 | 1.004 | 0.316 |  |
| Stressor (salinity) | 0.188 | 0.136 | 1.381 | 0.170 |  |
| Stressor (warming) | 0.141 | 0.101 | 1.395 | 0.164 |  |
| Microbe (prokaryotes) | 0.467 | 0.101 | 4.644 | <0.001 | *** |
| *Marginal R² = 0.196, Conditional R² = 0.440* | | | | | |

| ***(c) Compositional Shift Linear Mixed Model*** | | | | | |
| --- | --- | --- | --- | --- | --- |
| **Predictor** | **Estimate** | **SE** | **t** | **p** | **Sig.** |
| Modularity | 2.753 | 7.459 | 0.369 | 0.712 |  |
| Neg:Pos Cohesion | -3.240 | 5.303 | -0.611 | 0.542 |  |
| Network Richness | 0.010 | 0.009 | 1.074 | 0.284 |  |
| Broad Functional Redundancy | -0.024 | 0.027 | -0.919 | 0.359 |  |
| Mean Keystone Centrality | -1.931 | 1.044 | -1.850 | 0.065 | . |
| Keystone Retention Functions | -21.173 | 14.682 | -1.442 | 0.150 |  |
| Narrow Functional Redundancy | 1.932 | 2.023 | 0.955 | 0.340 |  |
| Stressor (salinity) | 10.300 | 3.906 | 2.637 | 0.009 | ** |
| Stressor (warming) | 4.086 | 2.467 | 1.656 | 0.099 |  |
| Microbe (prokaryotes) | -1.775 | 2.109 | -0.842 | 0.401 |  |
| *Marginal R² = 0.053, Conditional R² = 0.721* | | | | | |

| ***(d) Random Forest (10-fold CV)*** | |
| --- | --- |
| **Metric** | **Value** |
| AUC | 0.793 |
| Accuracy | 0.732 |
| Optimal mtry | 2 |
